# Haematological parameters and plasma levels of 8-iso-prostaglandin F2α in malaria-sickle cell co-morbidity: A cross sectional study

**DOI:** 10.1101/441998

**Authors:** Enoch Aninagyei, Emmanuel Doku Tetteh, Josephine Banini, Emmanuel Nani, Patrick Adu, Richard K. D. Ephraim, Alexander Egyir-Yawson, Desmond Omane Acheampong

**Author notes:** **Corresponding authors:** (AE), (AOD).

## Abstract

**Introduction:** Malaria and sickle cell disease (SCD) co-morbidity have previously been reported in Ghana. However, there is paucity of data on haematological profiles and oxidative stress in comorbidity states. This study identified novel inflammatory biomarkers associated with malaria in SCD and analyzed the levels of 8-iso-prostaglandin F2α oxidative stress biomarker in malaria-SCD co-morbidity in Ghanaian patients.

**Methods:** Blood (5ml) was collected from malaria patients into K_3_-EDTA tube. Malaria parasites speciation and quantification were then done according WHO guidelines. All eligible samples were assayed for haematological profile, sickle cell phenotyping, infectious markers (hepatitis B, hepatitis C, syphilis and HIV 1&2) and plasma levels of 8-epi-prostaglandin F2α..

**Results:** Prevalence of malaria in SCD (malaria-SCD) was 13.4% (45/335). Male: female ratio was 0.8:1 (X^2^=1.43, p=0.231). Mean ages for malaria in normal haemoglobin type (malaria-HbAA) and malaria-SCD were 12.79±4.91 and 11.56±3.65 years respectively (p=0.048). Geometric mean of parasite density was higher in malaria-HbAA (20394 parasites/μl vs. 9990 parasites/μl, p=0.001) whilst mean body temperature was higher in malaria-SCD (39.0±0.87°C vs. 37.9±1.15°C, p=0.001). Mean leukocytes, lymphocytes, eosinophils, monocytes, platelets and platelet indices values were significantly elevated in malaria-SCD. Significant reduction in RBC and RBC indices in malaria-SCD were also observed. Eosinophils-to-basophils ratio (EBR) and monocytes-to-basophils ratio (MBR) were novel cellular inflammatory biomarkers which could predict malaria in SCD. The sensitivities of cut-off values of EBR>14, MBR>22 and combined use of EBR>14 and MBR>22 were 79.55%, 84.09% and 91.11% respectively. Mean 8-iso-prostaglandin F2α was 338.1pg/ml in malaria-HbAA and 643.8pg/ml in malaria-SCD (p=0.001). 8-iso-prostaglandin F2α correlated with parasite density (r=0.787, p=0.001), temperature (r=0.566, p=0.001) and leucocytes (r=0.573, p=0.001) and negatively correlated with RBC (r=−0.476, p=0.003), haemoglobin (r=−0.851, p=0.001) and haematocrit (r=−0.735, p=0.001).

**Conclusion:** *Plasmodium falciparum* parasitaemia increases oxidative damage and causes derangement haematological parameters. Cut of values of EBR>14 and MBR>22 could predict malaria in SCD.

## Introduction

The asexual stages of *Plasmodium falciparum* are intra-erythrocytic thus inducing hematological alterations such as anemia, thrombocytopenia and neutrophilia [1–4]. Parasites density influence severity of the hematological changes. These changes depend on factors such as level of malaria endemicity, presence of haemoglobinopathies, nutritional status and level of malaria immunity [5, 6]. Malaria is meso-endemic in Ghana with recent nationwide prevalence of 43.4% [7]. Sickle cell disease (SCD) resulting from two haemoglobin S (HbS) haplotypes (HbSS) and heterozygote sickle cell phenotype resulting from one HbS haplotype and one haemoglobin C (HbC) haplotype (HbSC) is also prevalent in Ghana [8]. Sickle cell haplotype HbS leads to polymerization of deoxygenated sickle haemoglobin within inelastic red blood cells which cause occlusion of microvasculature, resulting in acute complications, chronic organ damage, high rate of morbidity and mortality [9]. These mechanisms have adverse effect on the quality and quantity of formed blood cells in affected individuals. SCD has been found to be associated with anemia, low RBC count, low packed cell volume (PCV), low mean cell volume (MCV), low mean cell hemoglobin (MCH) [10, 11] and leukocytosis [12]. The trends in the haematological profiles associated with SCD and malaria are similar. There is evidence of altered hematopoiesis which affect all the three hematological cell lines in SCD and malaria.

Oxidative stress is the overproduction of free radicals beyond the physiological detoxification ability of the body [13]. A principal consequence of Plasmodium infections is the development of anemia [14]. Malaria infections release reactive oxygen species (ROS) as a result of activation of the immune system of the body. Plasmodium infections and sickle cell disease are associated with oxidative stress due to production of ROS which result in haemoglobin degradation [15–17]. Oxidative stress is suspected to play a key role in disease pathogenesis, complications and mortality [18–19] consequently, a number of studies have focused on the measurement of oxidative stress, many of which are through specific biomarkers that indicate the oxidative damage [20]. Lipid, protein and DNA biomolecules are the main targets of free radicals that subsequently transformed into the reactive species reflecting oxidative stress in the corresponding molecules. Malondialdehyde (MDA), a product of lipid peroxidation, has been widely used as an indicator of oxidative stress. However, MDA measured by thiobarbituric acid assay overestimates actual MDA levels by more than 10-fold due probably to cross-reactivity with other aldehydes [21]. The isopentane, 8-iso-prostaglandin F2α (8-iso-PGF2α), is a more stable product of lipid peroxidation [22] and has been described at the gold standard biomolecule for assessing oxidative stress [23–24]. Measurement of 8-iso-PGF2α is a reliable tool for the identification of subjects with enhanced rates of lipid peroxidation [25].

In Africa, malaria and SCD are prevalent [26] and their co-morbidity have been reported in Ghana [27]. However, there is paucity of data on clinical manifestations, haematological profiles and degree of oxidative stress in SCD and malaria co-morbidities. Therefore, the aim of this study was to evaluate the variability of haematological parameters in malaria and SCD comorbidities, identify novel cellular inflammatory biomarkers for predicting malaria in SCD and levels of 8-iso-PGF2α oxidative stress biomarker in Ghanaian sickle cell patients infected with malaria parasites.

## Materials and methods

### Study site

This multi-center study took place in 3 district hospitals and 3 health centers in the Greater Accra region of Ghana. The hospitals (latitude, longitude) were Ga West Municipal Hospital, Amasaman (5.7020708, −0.2992889), Ashaiman Polyclinic (5.6856, −0.0398) and Ada East District Hospital (5.8956754, 0.5340865). The health centres (latitude, longitude) were Mayera Health Centre (5.720578, −0.2703561), Oduman Health Centre (5.64171, −0.3302) and Obom Health Centre (5.7361, −0.4395).

### Study design

A cross-sectional study conducted in malaria suspected patients from November 2017-August, 2018. The following clinical information; age, sex, temperature, body weight and clinical presentations was collected from each patient before sample collection. Maximum of three samples were collected each day from the study sites. 5ml of whole blood was collected from each consented patient. Specimens were transported from study sites to Ga West Municipal Hospital laboratory daily. Haematological profile was carried out on same day of sample reception. Blood films were prepared in triplicate. Screening for sickle cell and infectious makers were done, whole blood spun and plasma stored below −30°C. Haemolysate was prepared from concentrated red cell and kept frozen till ready for analysis.

### Study subjects, sample size and sample processing

Figure 1 details participant recruitment plan used for the present study. Patients recruited for the study were physician suspected malaria cases. Sample size was determined based on single population formula using confidence interval of 95% and estimated proportion of 1 in 4 malaria infections occurring in sickle cell disease. The sample size was estimated to be 289. Measurement of 8-epi-prostaglandin F2α biomarker was done in age and sex matched patients (normal controls n=40, malaria-HbAA n=40, SCD control n=40 and malaria-SCD n=40).

**Figure 1:**
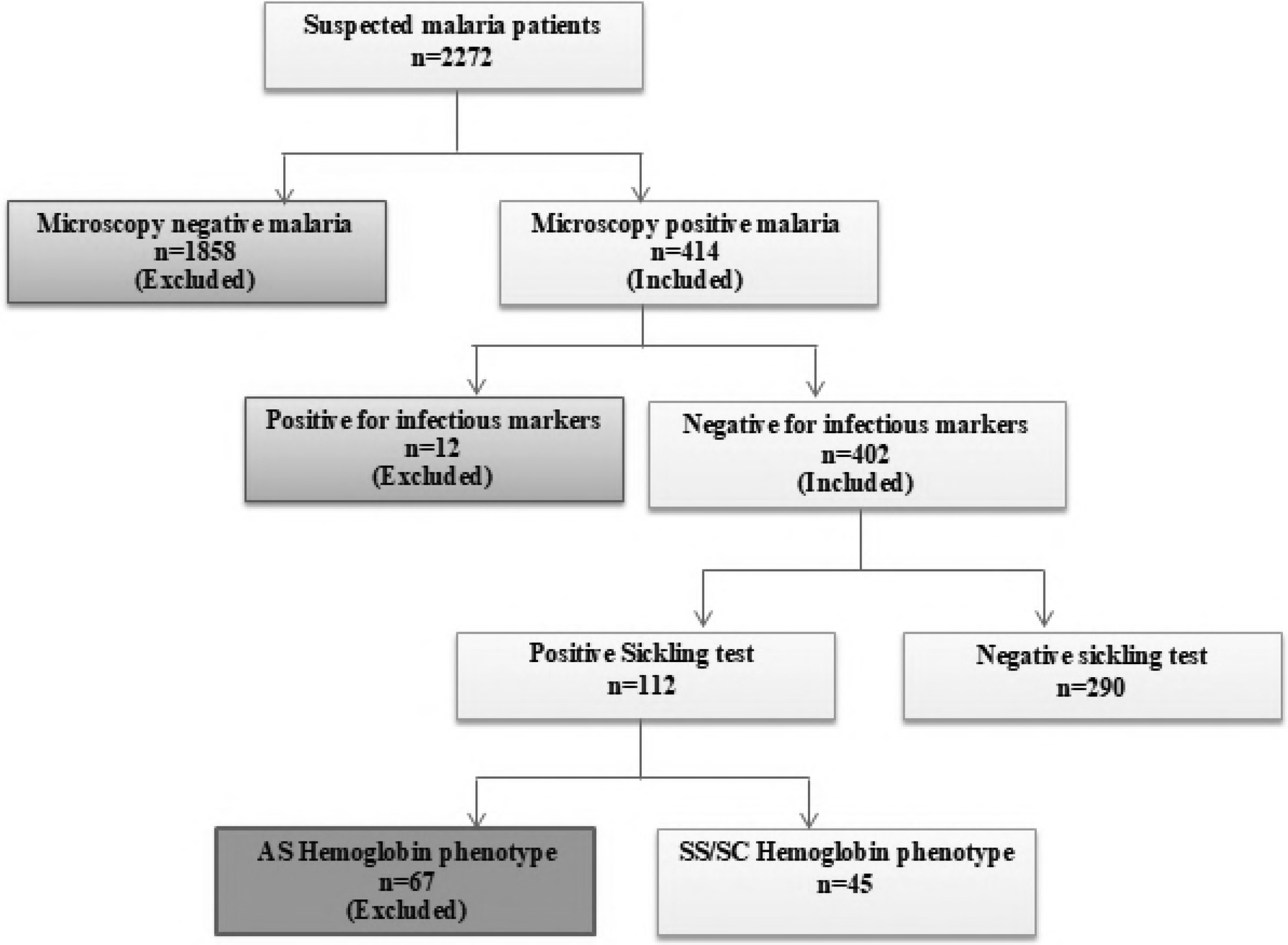
Flow chart for sample collection and analysis. Total of 2272 suspected malaria patients consented to participate in the study; microscopy did not detect malaria in 1858 (81.78%) of the patients while *Plasmodium falciparum* parasites were detected in 414 (18.22%) of the patients. Of the 414 malaria patients, 12 (2.9%) had hepatitis B surface antigen (5), hepatitis C antibody (1), Salmonella typhi IgM (8) and HIV I&II antibodies (3) detected in their plasma. 5 triple infections were identified. Of the 402 malaria mono-infected patients, 112 (27.8%) were sickle cell positive; 67(60.0%) and 45(40.0%) were malaria in haemoglobin AS and haemoglobin SS/SC respectively. Prevalence of SCD and sickle cell trait (SCT) with malaria were 11.2% (45/402) and 16.6% (67/402) respectively. Excluding malaria in SCT from the study, the prevalence of malaria in SCD was computed to be 13.4% (45/335).

### Inclusion and exclusion criteria

Patients included in the study were microscopy diagnosed malaria patients, aged 0-20 years, who consented or whose parents consented to be part of the study. Individuals who were known SCD patients or visited the health centre on account of sickle cell crisis and malaria patients co-infected with hepatitis B virus, hepatitis C virus, syphilis and HIV 1&2 were excluded. Also samples with malaria and sickle cell trait were excluded.

### Laboratory analysis

#### Malaria detection and quantification

Thick and thin blood film was done for each specimen, in triplicate, on the same glass slide. The dried thin film was fixed in absolute methanol briefly, air dried and stained with 10% Giemsa. The dried smear was then examined for presence of Plasmodium parasites. The parasites were subsequently identified to the species level and quantified per μl of blood and percentage RBC infected according WHO guidelines. Parasites were quantified per 200 WBCs counted using the patients’ total WBC per μL of blood. A total of 500 WBCs were counted in negative infections [28]. Each slide was double checked by a blinded certified malaria microscopist and in cases of discordant results; a third opinion was final.

#### Infectious marker screening

The specimens were screened for hepatitis B virus, hepatitis C virus, syphilis and HIV I&II pathogens to eliminate any possible effect on the haematological parameters. The microbiological screening was done with rapid immunochromatographic test kits. HIVI&II and syphilis were screened with First Response^®^ test kit (Premier Medical Corporation Ltd, India) whilst the hepatitis B and C were screened with FaStep Rapid Diagnostic Test (Houston, USA).

#### Haematological profiling

Haematological profiling was done using Urit 5200 (China) fully automated haematology analyzer. The 5-part differential analyzer works on the principle of laser beam multidimensional cell classification flow cytometry for white cell differentiation, white and red blood cell estimation. Platelets were counted by optical and electrical impedance principles and haemoglobin concentration was measured by cyanide-free colorimetric method. All other parameters were calculated.

#### Determination of leukocyte ratio cut-off values

The 95% confidence interval (CI) was determined for each mean of the leukocyte ratios recorded in malaria-SCD. The upper or the lower value of the CI that majority of the individual values fell was taken as the cut-off value. The cut-off values were determined on ratio by ratio basis.

#### Sickle cell screening and phenotyping

Sickle cell screening was done by the sodium metabisulphite reduction method as previously described by Antwi-Baffour [29] Sickle cell phenotyping was done as described by Cheesborough [30]. Haemolysate was separated using electrophoresis in alkaline medium (pH 8.6). Haemoglobin phenotyping was done alongside pooled HbA, HbS, HbC and HbF controls. Electrical voltage of 250 V and current of 50 mA were employed to obtain complete separation of haemoglobin variants for a maximum of 30 min. Results were read immediately against the controls.

#### Sandwich-ELISA for 8-epi-prostaglandin F2α levels

Reagents and consumables for 8-epi-prostaglandin F2alpha was obtained from SunLong Biotech (Hangzhou, China, Catalogue Number: SL0035Hu). Measurement of 8-epi-prostaglandin F2alpha was done according to manufacturer’s protocol, with these modifications; incubation of pre-diluted samples with coated anti-8-epi-prostaglandin F2α and after addition of Horseradish Peroxidase-conjugated antibody specific for human 8-epi-prostaglandin F2alpha was done for 45 minutes at room temperature. Again, the chromogen solution was incubated for 30 minutes at room temperature after dispensing into the microelisa wells. The optical density (OD) was measured by Mindray MR-96A ELISA plate reader (Shenzhen, China) at a wavelength of 450 nm. The concentration of 8-epi-prostaglandin F2α was obtained by comparing the OD of the samples to the standard curve.

### Statistical analysis

Raw data were entered into Microsoft Excel 2010. Statistical analyses were done with Stata 15.0 statistical software (Stata Corp LLC, USA). Pearson Chi square was used to determine differences in categorical data whilst differences in parametric variables were determined by t-test. To compare more than two groups, one-way ANOVA was used with multiple comparison undertaken using Tukey post hoc analysis. Correlation between variables was determined by Pearson correlation test. P-value <0.05 was considered statistically significant. Receiver operating characteristic (ROC) curve was used to estimate the sensitivity, specificity and. predictive values of eosinophils-to-basophils and monocytes-to-basophils ratios to predict malaria in sickle cell disease.

## Results

### Demographic, temperature and parasitemia in malaria and sickle cell co-morbidity

Patients with normal haemoglobin but infected with malaria (malaria-HbAA) formed the majority (86.6%; 290/335) of participants compared to malaria-SCD (13.43%; 45/335). Majority of the participants were females (55.8% females vs. 44.2% males). The mean age of malaria-HbAA was significantly higher that of malaria-SCD patients (12.79±4.91 years vs 11.56±3.65 years; p = 0.048). Body temperature was significantly higher in malaria-SCD than malaria-HbAA (37.9±1.15 vs. 39.0±0.87; t=6.86, p=0.001). The geometric mean of malaria parasite density was higher in malaria-HbAA (20394 parasites/μL, IQR 9519-51093) than malaria-SCD (9990 parasites/μL, IQR 6329-16945) (table 1).

**Table 1:** Demographic, temperature and parasitaemia of the patients.

| Variables | Malaria-HbAA<br>(n=290) | Malaria –SCD<br>(n=45) | Statistic | p-value |
| --- | --- | --- | --- | --- |
| Age (mean±SD) | 12.79±4.91 | 11.56±3.65 | t=2.01 | 0.048<sup>a</sup> |
| Males, n (%) | 127 (43.80) | 21 (46.70) | X<sup>2</sup>=1.43 | 0.231<sup>b</sup> |
| Females, n (%) | 163 (56.20) | 24 (53.30) |  |  |
| Body temp range (mean ±SD) | 37.9±1.15 | 39.0±0.87 | t=6.89 | <0.001<sup>a</sup> |
| Parasite density range | 223-620586 | 2492-112452 |  |  |
| GM of Parasite density | 20394 | 9990 | t=7.43 | <0.001 |
| Interquartile range | 9519-51093 | 6329-16945 |  |  |

<sup>a</sup> p-value determined by t-test, <sup>b</sup> p-value determined by Pearson Chi square, GM-Geometric mean

### Haematological parameters in malaria and sickle cell co-morbidity

The haematological variables of the participants were also compared (table 2). Whereas TWBC (12.32±2.77 vs., 6.68±2.42 p=0.001), %lymphocytes (36.23±8.44 vs. 28.53±18.22, p=0.001), %eosinophils (4.77±0.99 vs. 2.19±1.79, p=0.001) and %monocytes (7.32±1.58 vs. 5.92±3.30, p=0.001) were significantly higher in malaria-SCD, %neutrophil (62.1 ±20.1 vs. 50.44±8.65, p=0.001) and %basophils (0.45± 0.24 vs. 0.32±0.07, p=0.001) were significantly higher in malaria-HbAA participants. Also, RBC count (4.22±0.78 vs. 3.87±0.69, p=0.001), haemoglobin (9.19±1.06 vs. 10.83±2.11, p=0.001), haematocrit (27.34±2.79 vs. 31.84±6.07, p=0.001), MCV (71.33 ±7.62 vs. 76.07±10.53, p=0.001) and MCH (24.53±4.09 vs. 25.89±3.78, p=0.041) were significantly lower in malaria-SCD than malaria-HbAA. Although MCHC and RDW_CV differed among the participants, the differences did not reach statistical significance. Moreover, platelets (190.0± 55.30 vs. 138.71±87.70, p=0.001), MPV (10.78±1.28 vs. 9.91±1.40, p=0.001), PDW (13.33±1.89 vs. 12.66±2.48, p=0.038), P_LCR (30.21±7.39 vs. 25.11±8.06, p=0.001) were significantly higher in malaria-SCD than malaria-HbAA.

**Table 2:**
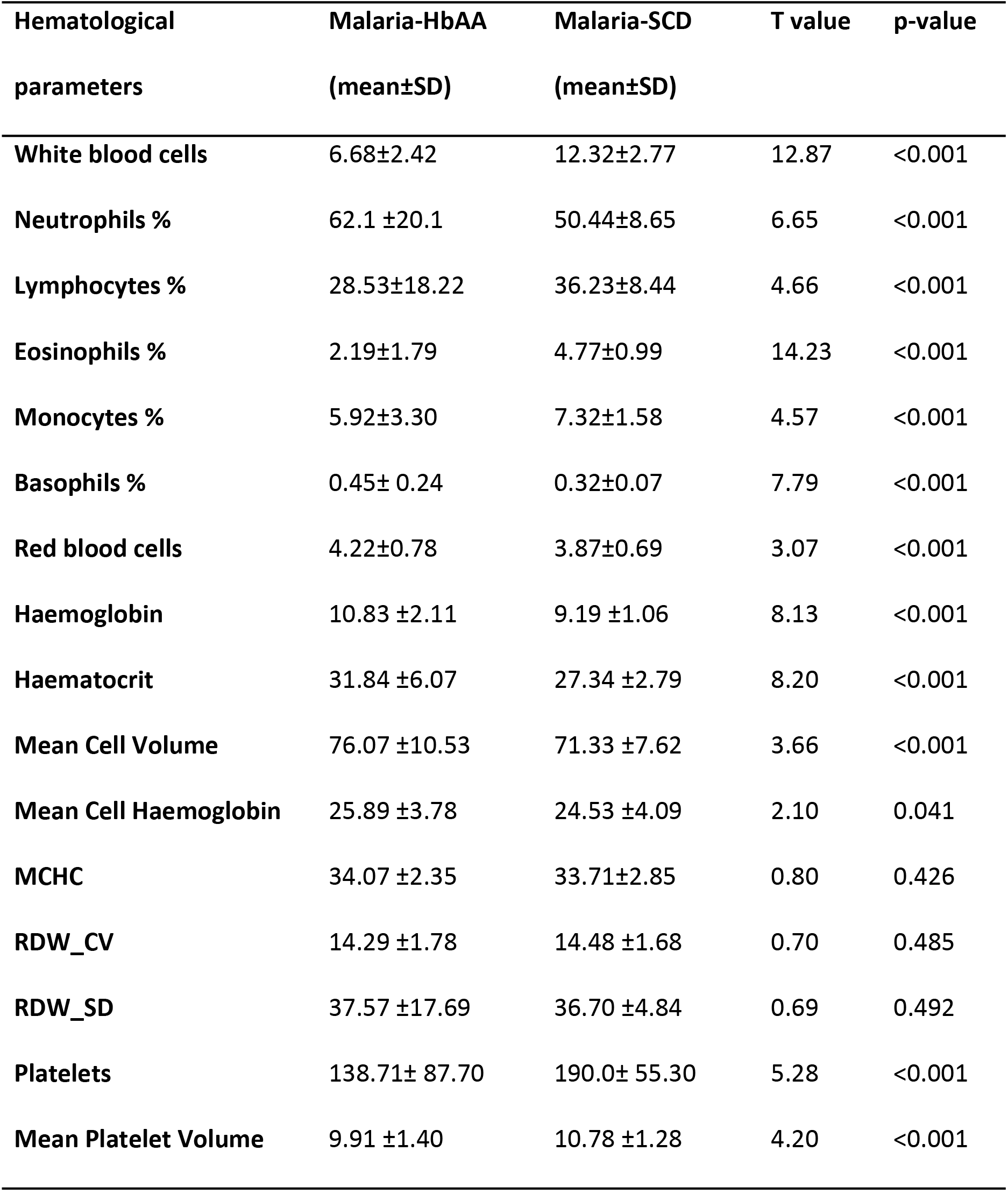
Hematological parameters of malaria and sickle cell co-morbidity.

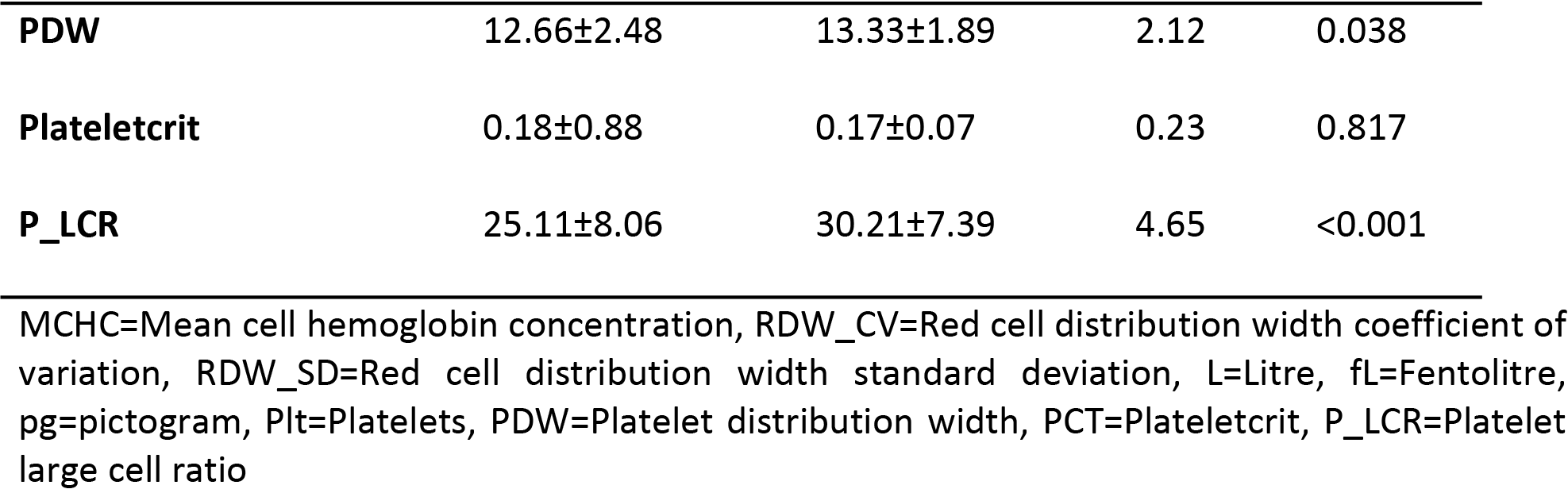

### Variations in leukocyte ratios in malaria and sickle cell co-morbidities

The mean leukocytes ratios observed in malaria-HbAA and malaria-SCD were significantly different from each other (table 3). Lymphocytes-to-basophils ratio (76.10±66.33 vs. 125.19±59.30, p=0.001), eosinophils-to-monocytes ratio (0.43±0.65 vs. 0.68±0 .21, p=0.001), eosinophils-to-basophils ratio (4.62±2.52 vs. 16.05±5.64, p=0.001), monocytes-to-basophils ratio (12.85±2.79 vs. 23.40±3.04, p=0.001) and platelets-to-neutrophils ratio (2.85±3.23 vs. 3.82± 1.04, p=0.001) were significantly higher in malaria-SCD. However, neutrophils-to-lymphocytes ratio (3.82±3.86 vs. 1.51±0.55, p=0.001), neutrophils-to-eosinophils ratio (13.08±5.87 vs. 11.30±4.22, p=0.001), neutrophils-to-monocytes ratio (16.97±25.18 vs. 7.25± 2.35, p=0.001), lymphocytes-to-eosinophils ratio (55.85±9.43 vs. 7.88±2.38, p=0.001), lymphocytes-to-monocytes ratio (8.32±8.48 vs. 5.33± 2.27, p=0.009) and platelets-to-lymphocytes ratio (7.16±6.80 vs. 5.63± 2.57, p=0.006) were significantly lower in malaria-SCD.

**Table 3:**
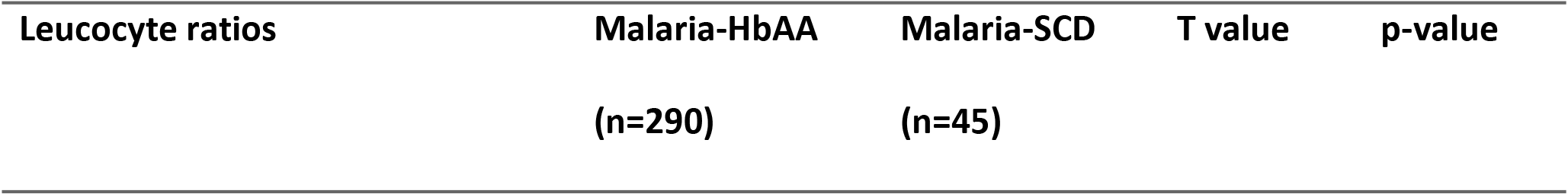
Leukocyte ratios among malaria and malaria-sickle cell co-morbidities.

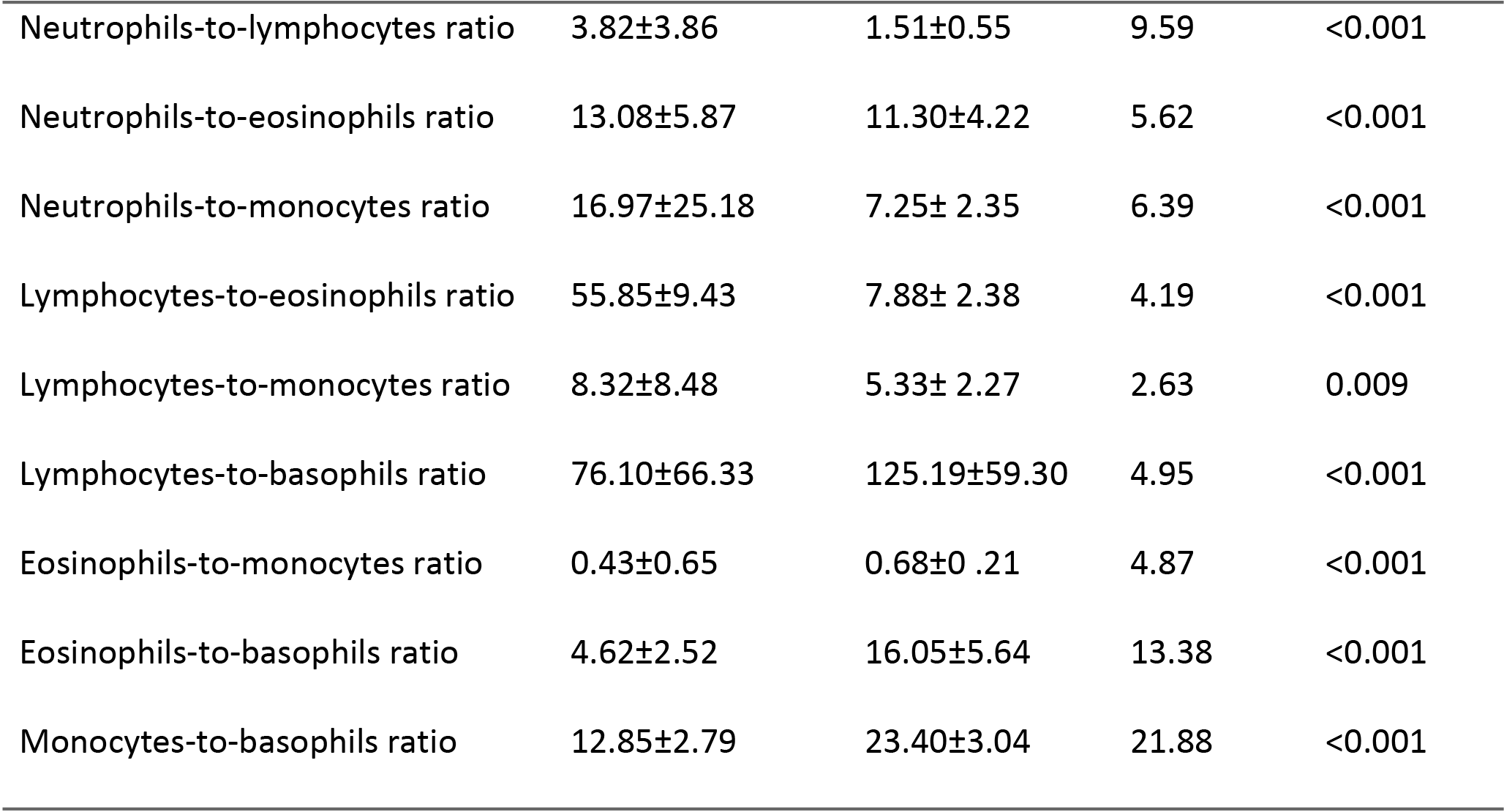

### Predictive novel cellular inflammatory biomarkers in malaria in SCD

In table 4 the leucocyte ratios were further explored for their ability to predict malaria-SCD comorbidity by means of receiver operating characteristic (ROC). Neutrophils-to-monocytes ratio was the most sensitive (93.33%, 95% CI: 81.73-98.60) but only 56.90% specific (95% CI: 50.9862.67) and had very low positive predictive value (PPV) (25.15%, 95% CI 22.37-28.15). Lymphocytes-to-eosinophils ratio and lymphocytes-to-monocytes ratio were 86.67% and 80.00% sensitive respectively but not very specific (65.17% and 35.86% respectively) and had very low PPV (27.86% and 16.22% respectively). The sensitivity, PPV and ROC of lymphocytes-to-basophils ratio, neutrophils-to-eosinophils ratio and neutrophils-to-lymphocytes were comparatively low. Eosinophils-to-basophils ratio (EBR) and monocytes-to-basophils ratio (MBR) had relatively high predictive values. The cut-off values for EBR>14 and MBR>22 associated with malaria-SCD. The sensitivity, specificity, positive predictive value (PPV), negative predictive value (NPV) and ROC of EBR>14 were 79.55% (95% CI: 64.70-90.20), 97.11% (95% CI: 94.75-98.61), 77.78% (95% CI: 65.11-86.78), 97.39 % (95% CI: 95.42-98.53) and 88.33% (95% CI: 79.73-94.40). The values obtained for MBR>22 in predicting malaria in SCD were 84.09 % sensitive (95% CI: 69.93-93.36), 97.69 % specific (95% CI: 95.50-99.00), 82.22% PPV (95% CI 69.73-90.28), 97.97% NPV (95% CI: 96.07-98.96) and ROC value 90.89% (95% CI: 82.72-96.18). When the combined performance of both EBR>14 and MBR>22 were analyzed, the obtained values were higher than using either ERB>14 or MBR>20 alone. The following indices were thus obtained for EBR>14-MBR>22; 91.11% sensitivity (95% CI: 78.78-97.52), 98.55% specificity (95% CI: 96.66-99.53), 89.13% PPV (95% CI: 77.37-95.16), 98.84% NPV (95% CI: 97.10-99.54) and ROC 94.83% (95% CI 87.72-98.52). EBR and MBR were novel inflammatory markers found to be significantly associated with malaria-SCD.

**Table 4:**
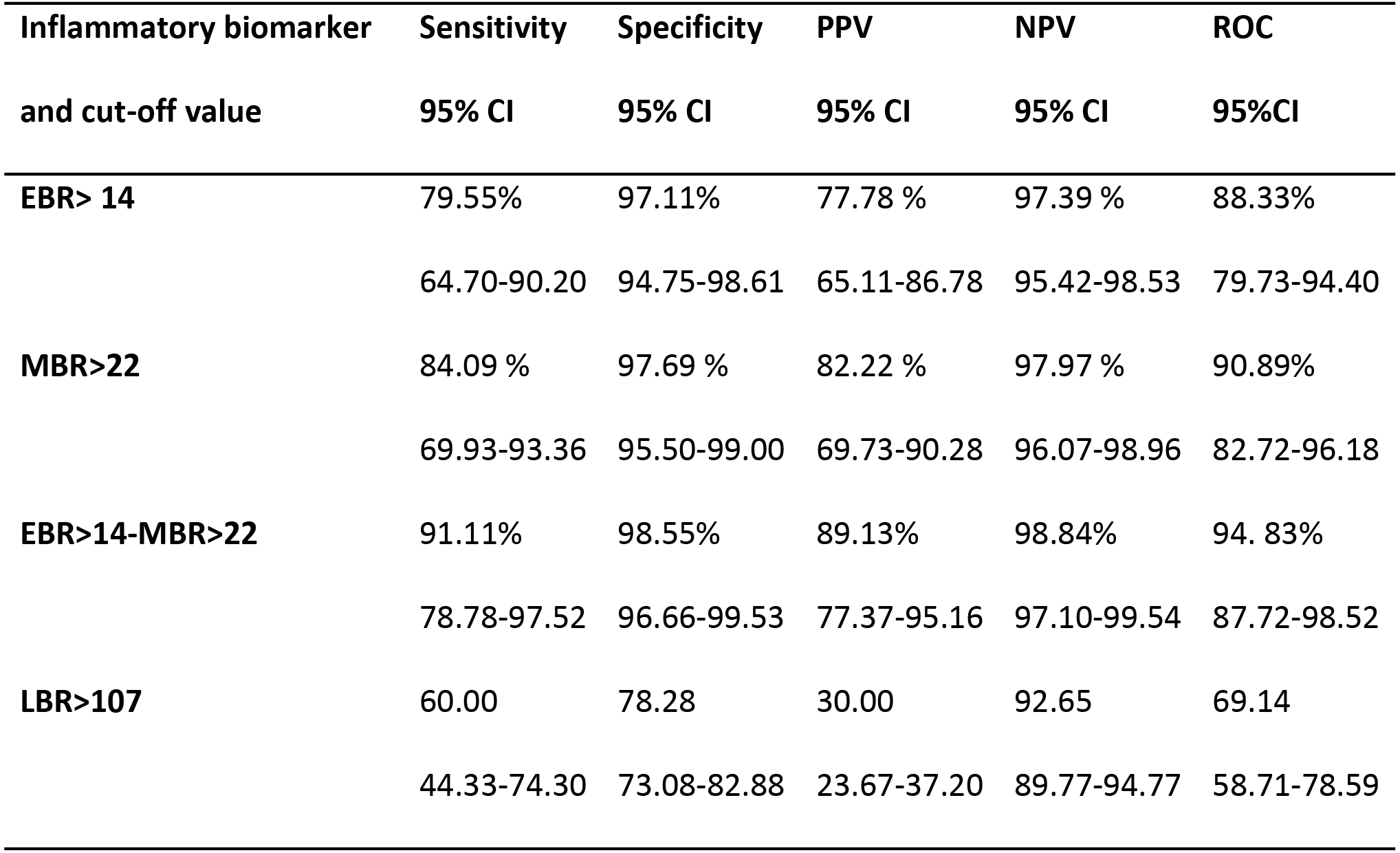
Novel leukocyte ratios associated with malaria-SCD.

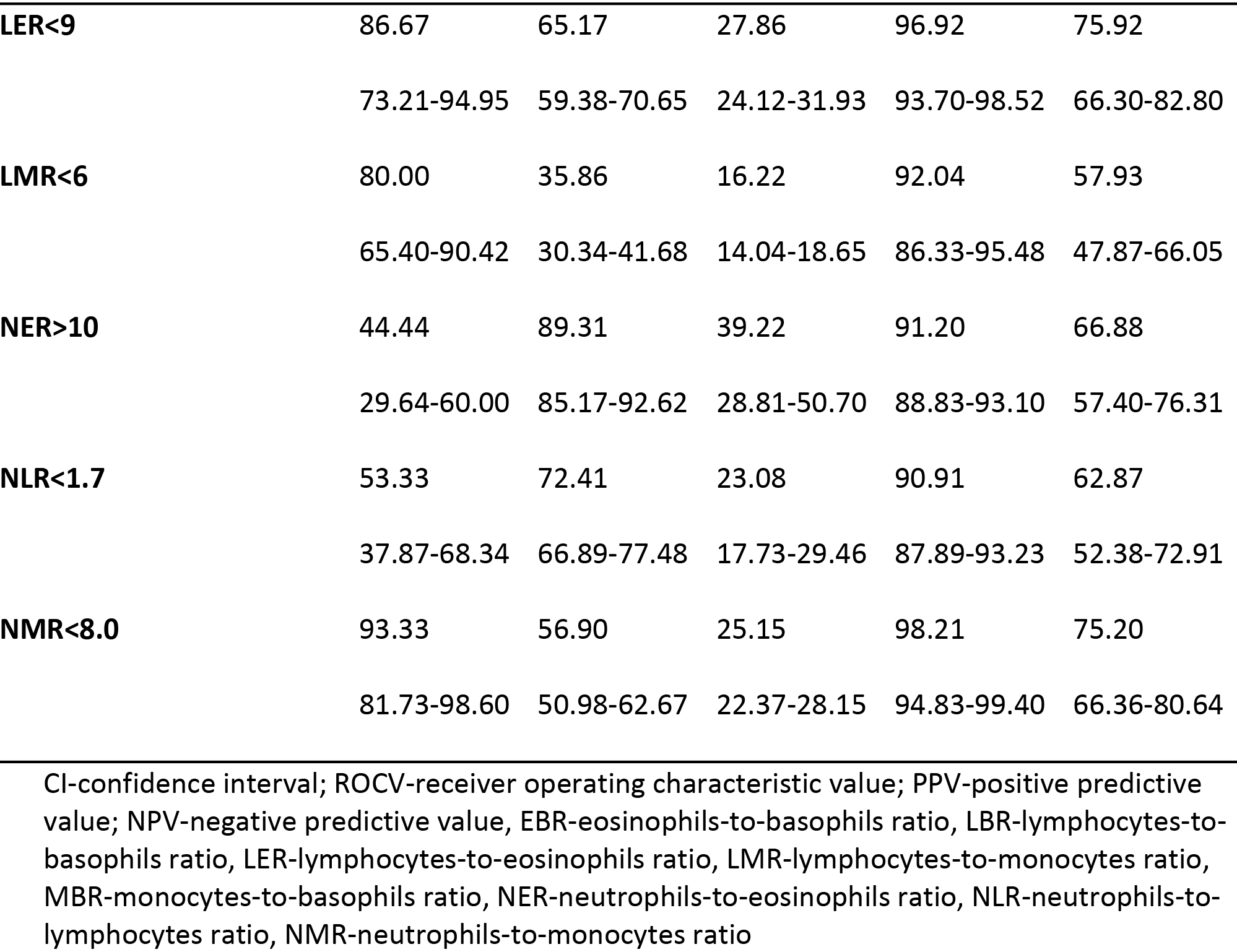

### 8-iso-prostaglandin F2α levels and its correlates

The oxidative stress associated with malaria infection and/or SCD inheritance were also assessed through estimation of 8-iso-prostaglandin F2α (table 5). The mean levels of 8-isoprostaglandin F2α oxidative stress biomarker was significantly lower in control (no malaria-SCD negative participants) compared to all other groups [84.1pg/ml vs 129.1pg/ml (SCD patients) vs 338.1pg/ml (malaria-HbAA) vs 643.8pg/ml (malaria-SCD); p = 0.001]. However, Tukey post hoc analysis indicated non-significance difference between malaria-SCD and malaria-HbAA (t=1.13, p=0.792). But pairwise post hoc analysis in the other groups were significant [(Malaria-SCD vs. normal control, t=41.12, p=0.001), (Malaria-SCD vs. SCD, t=36.5, p=0.001), (Malaria-HbAA vs.normal control, t=40.0, p=0.001), (Malaria-HbAA vs. SCD, t=35.38, p=0.001) and (SCD vs. normal control, t=44.95, p=0.001)]. Moreover, whereas 8-iso-prostaglandin F2α was significantly positively correlated with parasite density (r=0.787, p=0.001), temperature (r=0.566, p=0.001) and total WBC (r=0.573, p=0.001), it was inversely related to RBC (r=−0.476, p=0.003), haemoglobin concentration (r=−0.851, p=0.001) and haematocrit (r=−0.735, p=0.001).

**Table 5:** Analysis of 8-iso-prostaglandin F2α and its correlation with haematological and other parameters.

| Variable | Control | SCD patients | Malaria-HbAA | Malaria-SCD | p-value |
| --- | --- | --- | --- | --- | --- |
| Age (years) | 13.875(3.8) | 14.125(3.7) | 13.525(4.2) | 13.575(3.7) | 0.956<sup>a</sup> |
| **Gender** |  |  |  |  |  |
| Male | 17(42.50) | 20(50.00) | 16(40.00) | 18(45.00) | 0.675<sup>b</sup> |
| Female | 23(57.50) | 20(50.00) | 24(60.00) | 22(55.00) |  |
| 8-iso-PGF$2\alpha$ | 84.1(16.3) | 129.1(18.8) | 338.1(50.2) | 643.8(54.54) | <0.001<sup>a</sup> |
| **8-iso-PGF$2\alpha$ levels correlates with significant variables** |  |  |  | **r** | **p-value** |
| Parasite density |  |  |  | 0.787 | <0.001 |
| Body temperature |  |  |  | 0.566 | <0.001 |
| Red blood cells |  |  |  | -0.476 | 0.003 |
| Haemoglobin |  |  |  | -0.851 | <0.001 |
| Haematocrit |  |  |  | -0.735 | <0.001 |
| Total WBC |  |  |  | 0.573 | <0.001 |

| 8-iso-prostaglandin F2α levels | Tukey | |
| --- | --- | --- |
| Tukey pairwise analysis | t | p-value |
| Malaria-SCD vs. malaria-HbAA | 1.13 | 0.792 |
| Malaria-SCD vs. normal control | 41.12 | <0.001 |
| Malaria-SCD vs. SCD | 36.50 | <0.001 |
| Malaria-HbAA vs. normal control | 40.00 | <0.001 |
| Malaria-HbAA vs. SCD | 35.38 | <0.001 |
| SCD vs. normal control | 4.62 | <0.001 |

<sup>a</sup>one-way-ANOVA; <sup>b</sup>Pearson chi square; <sup>c</sup>post-hoc analysis; abbreviations: 8-iso-PGF2α -8-iso-prostaglandin F2α ; SCD-sickle cell disease; Malaria-SCD-malaria-sickle cell disease co-morbidity

## Discussion

In sub-Saharan Africa where inheritance of SCD is high and malaria infection is endemic, malaria and SCD comorbidity is prevalent in the general population. Malaria has been demonstrated to induce oxidant stress due to parasite multiplication as well as host immune response to the parasite [31]. In this study we show that inheritance of SCD alone or P. falciparum infection induces significant elevations in oxidant stress marker 8-iso-prostaglandin F2α. Additionally, P. falciparum infection and SCD co-morbidity leads to synergistic increase of this oxidant stress biomarker in the peripheral blood of these patients.

The utility of 8-iso-prostaglandin F2α as oxidative stress biomarker indicated significant increases in malaria-HbAA, malaria-SCD and sickle cell patients compared to control subjects. 8-iso-prostaglandin F2α oxidative stress biomarker levels increased by 1.53, 4.0 and 7.6 folds in SCD, malaria-HbAA and malaria-SCD respectively. This finding is suggestive that 8-isoprostaglandin F2α may be a useful oxidative stress biomarker in malaria and sickle cell patients. Interestingly levels of 8-iso-prostaglandin F2α in malaria-SCD comorbidity significantly correlated positively with parasite density, total WBC and body temperature and negatively with RBC, haemoglobin and haematocrit. As it has been demonstrated that increasing parasitemia leads to increased red cell destruction, elevated body temperature of patients and subsequent oxidant stress, the relationship between 8-iso-prostaglandin F2α and P. falciparum parasitemia reported herein is not surprising. The mean 8-iso-prostaglandin F2α in malaria-SCD patients was twice the mean 8-iso-prostaglandin F2α in malaria-HbAA patients. The observed differences could be attributed to cumulative effect of malaria infection in SCD. Previous study in Kenya found SCD to increase the severity of malaria [32]. This study could link this finding to excessive lipid peroxidation with the consequent elevation of 8-iso-prostaglandin F2α. There is limited data on the use of 8-iso-prostaglandin F2α oxidative to assess oxidative stress in SCD even though previous works have assessed oxidative stress using nitric Oxide (NO), superoxide, peroxide, hydroxyl radicals [33, 34] and malondialdehyde (MDA) [33, 35–37]. The use 8-isoprostaglandin F2α to assess oxidative stress in malaria is not popular, though 8-isoprostaglandin F2α is currently the gold standard for the assessment of oxidative stress in disease conditions [23, 24]. Oxidative stress due to reactive oxygen species (ROS) activity on MDA has been implicated previously in the pathogenesis and complications in malaria. It has been established that Falciparum infected human RBCs are under constant oxidative stress [38] due to generation of ROS within erythrocytes infected cells and also from immune activation [39, 40].

Previous study done in Ghana in patients below 20 years, reported that WBC and lymphocytes in malaria patients were lower compared to control subjects. The other leucocytes sub-types were not significantly different from control subjects [41]. However, the current study observed significantly high leucocytes and leucocytes sub-types derangements in malaria-SCD except neutrophils and basophils. Presentations of observed symptoms in malaria-SCD could be related to the actions of pro-inflammatory cytokines with mononuclear cells having been implicated as a key player [42]. This study found two novel cellular inflammatory biomarkers, namely eosinophils-to-basophils ratio (EBR) and monocytes-to-basophils ratio (MBR), as being associated with malaria-SCD. EBR>14 and MBR>22 and combination of the two, EBR>14-MBR>22, could be used to predict malaria in sickle cell disease. The sensitivity of EBR>14-MBR>22 was 11.56% and 7.02% higher than EBR>14 and MBR>22 when compared individually. The specificity and negative predictive values of the novel biomarkers were greater than 90%; this makes them very specific in excluding malaria in sickle cell disease. The ROCV obtained makes the diagnostic use of EBR>14-MBR>22 better than EBR>14 and MBR>22. As leukocyte ratios have been widely suggested to predict both communicable and non-communicable diseases [43–45], our findings are suggestive of potential roles in P. falciparum pathophysiology.

In malaria-SCD, haemoglobin, hematocrit, mean cell haemoglobin and mean cell volume were significantly reduced in a similar fashion as seen in microcytic and hypochromic anaemia. The reductions in RBC and red cell indices associated with malaria-SCD could probably be due to cumulative effect of increased rate of haemolysis during oxygenation and deoxygenation process, reduced response to erythropoietin secretion in sickle cell anemia together with acute malaria infections [46]. Significant elevations were observed in platelets, mean platelet volume, platelet distribution width and plateletcrit in malaria-SCD patients. Relative thrombocytopenia was seen in the malaria-HbAA patients. Malaria-related case vs. control thrombocytopenia has been reported in several studies [6, 47–48]. Elevation in platelets and platelet indices suggests efficient haemostasis in malaria-SCD than malaria infections in the absence of SCD.

## Conclusion

EBR>14 and MBR>22 were novel cellular inflammatory biomarkers found to be associated with malaria-SCD and can possibly be employed in the diagnosis of this co-morbidity. Additionally, malaria-SCD levels of 8-iso-prostaglandin F2α oxidative stress biomarker was twice as observed in malaria-HbAA.

## List of abbreviations

ANOVA-one way analysis of variance; EBR-eosinophils-to-basophils ratio; ELISA-enzyme lined immuno-sorbent assay; EMR-eosinophils-to-monocytes ratio; HbS-Sickle cell hemoglobin; HbSS-two HbS haplotypes; LBR-lymphocytes-to-basophils ratio; LER-lymphocytes-to-eosinophils ratio; LMR-lymphocytes-to-monocytes ratio (LMR); malaria-HbAA-malaria in normal hemoglobin; malaria-SCD-malaria in sickle cell disease; MDA-malondialdehyde; MBR-monocytes-to-basophils ratio; MCH-mean cell hemoglobin; MCV-mean cell volume; NBR-neutrophils-to-basophils ratio; NER-neutrophils-to-eosinophils ratio; NLR-neutrophils-to-lymphocytes ratio; NMR-neutrophils-to-monocytes ratio; OD-optical density; PNR-platelet-to-neutrophils ratio; ROC-receiver operating characteristic; SCD-Sickle cell disease; SCT-sickle cell trait; SSA-Sub-Saharan Africa; TBAA-thiobarbituric acid assay

## Ethical approval

Ethical approval for this study was granted by Ghana Health Service Ethics Review Committee (Approval No: GHS-ERC002/03/18). Participant consent was sought for participant.

## Raw data

All relevant data are within the paper.

## Acknowledgements

We acknowledge Michael Ankwadah, Michael Gyimah, Sedem Bokor, Bridgette Tevi and Alex Nyarko for the role they played in recruiting the patients, collection of clinical data and taking specimens for the study. We are also grateful to Nicholas Sowa for his immense contributions during the laboratory measurement of the 8-iso-prostaglandin F2α oxidative stress biomarker.

## Authors′ contributions

EA, PA, DOA, RKDE conceptualized, designed and coordinated the study. EA, EDT, PA performed the statistical analysis and JB, EN participated in the sample collection and processing. AE, PA drafted the manuscript, manuscript proofread by RKDE, AE-Y which was later approved by all authors

